# MyeliMAP: Studying Oligodendrocyte Function in Brain Networks

**DOI:** 10.1101/2025.11.14.688409

**Authors:** Karan Ahuja, Blandine F. Clément, Giulia Amos, Joël Küchler, Keimpe Wierda, Yoke Chin Chai, Lieve Moons, Catherine Verfaillie

## Abstract

Oligodendrocytes are the myelinating glia of the central nervous system (CNS), essential for rapid signal propagation, metabolic support, and neuronal health. While rodent-based cultures and organoid systems have provided insights into oligodendrocyte biology, they fall short of capturing human-specific features of myelination or integrating structural and functional readouts. Here, we present MyeliMAP (<u>Myeli</u>nation <u>MAP</u>ping), a human pluripotent stem cell (hPSC) -derived microphysiological and electrophysiological platform that enables robust modeling of CNS myelination. The system combines inducible hPSC-derived neurons and oligodendrocytes in a custom-engineered microfluidic microstructure designed to mimic the developing brain microenvironment, promoting spatially organized axon-glia interactions and controlled myelin sheath formation. Within six weeks, we demonstrate myelin formation and maturation by immunofluorescence and ultrastructural validation using transmission electron microscopy (TEM), confirming compact multilayered wrapping of human axons. Importantly, the microstructure is directly integrated with a high-density microelectrode array (HD-MEA), enabling real-time, long-term functional assessment of network activity and myelin-dependent changes in signal conduction. This allowed us to demonstrate that oligodendrocyte-based myelinated neurons display enhanced conduction velocity of action potentials compared to neuron monocultures. Moreover, the presence of oligodendrocytes stabilized the temporal neuronal network activity by reducing variability in firing patterns and enhancing synchrony across the culture. This dual structure-function approach surpasses static end-point analyses by coupling morphological validation with dynamic, quantitative measurements of maturing circuit physiology. MyeliMAP provides a reproducible, human-relevant platform to dissect neuron-glia interactions and accelerate discovery of remyelination-promoting strategies for CNS disease.

## Introduction

Oligodendrocytes are the myelinating glial cells of the central nervous system (CNS), essential for rapid action potential propagation and long-term neuronal health. Beyond providing electrical insulation through compact myelin, these cells contribute to axonal metabolic support, ion homeostasis, and synaptic regulation^1^. Disruption of oligodendrocyte function or myelin integrity is a hallmark of numerous neurological conditions, ranging from demyelinating diseases such as multiple sclerosis (MS) to neurodegenerative disorders including Alzheimer’s disease (AD)^2^. Understanding how oligodendrocytes interact with neurons and other glial cell types is therefore critical for developing strategies to preserve or restore myelin in disease.

Advances in human pluripotent stem cell (hPSC) technology have transformed our ability to generate human oligodendrocytes *in vitro*, enabling the creation of species-relevant models to investigate myelination and its dysregulation in disease. Traditional *in vitro* systems, such as rodent primary cultures, immortalized cell lines, or even human neuron-oligodendrocyte cocultures, have been instrumental in uncovering fundamental aspects of oligodendrocyte development^3,4^. However, these platforms fall short in recapitulating the cellular diversity, human-specific molecular signatures, and three-dimensional (3D) microenvironment of the CNS. More advanced models, including brain organoids and compartmentalized microfluidic devices, now provide 3D architecture or spatially controlled axon-oligodendrocyte interactions that permit myelin formation^5–7^. Yet, most of these systems remain reliant on static end-point analyses, such as myelin basic protein (MBP) immunostaining or electron microscopy, offering only a snapshot of myelination without capturing its dynamic or functional impact on axonal conduction.

One underexplored aspect of *in vitro* myelination models is the contribution of oligodendrocyte progenitor cells (OPCs) to neuronal development and, ultimately, myelin formation. Beyond their role as precursors of mature oligodendrocytes, OPCs form synaptic contacts with neurons, sense network activity, and secrete trophic factors^8,9^. These interactions position OPCs as active modulators of neuronal excitability and circuit maturation, which in turn affect the capacity of neurons to undergo functional myelination^10^. However, the temporal link between OPC-mediated neuronal maturation and subsequent myelin formation remains poorly defined in human-based systems. Functional assessment of this stepwise contribution—first to neuronal network stabilization, and later to myelin-dependent conduction—offers critical insight into how oligodendrocytes support circuit function and why these processes are disrupted in demyelinating diseases^11^.

Among human brain models used to study neuron-oligodendrocyte codifferentiation and maturation, inducible *NGN2* neurons (iNGN2 neurons) offer a rapid, reproducible, and subtype-specific source of excitatory human neurons, making them highly suitable for standardized, high-throughput investigations of neuron-glia interactions and drug discovery.^12,13^. Despite their broad application in neuronal specification, maturation, and drug screening, iNGN2 neurons have yet to demonstrate consistent and robust myelination. Although partial axon-oligodendrocyte associations have been reported^14,15^, definitive ultrastructural evidence of compact myelin and node of Ranvier formation remains absent.

Here, we present <u>Myeli</u>nation <u>MAP</u>ping (MyeliMAP), a microphysiological platform integrating electrophysiological assessment, designed to capture both the structural and functional hallmarks of human CNS myelination. Using iNGN2 neurons, we show for the first time that these cells can be myelinated when cocultured with hPSC-derived inducible *SOX10* OPCs (iSOX10 OPCs)^16^ in the MyeliMAP platform. This creates a human-relevant system that links structural and functional readouts of myelination within a short experimental timeline. We show that cocultured iNGN2 neurons and iSOX10 OPCs in MyeliMAP platform form functional networks that reproduce key steps of human CNS myelination. Neuronal extensions (neurofilament H (NF) positive) ensheathment by oligodendrocytes (Myelin Basic Protein (MBP) positive) and paranodal organization (Contactin-Associated Protein (CASPR) positive) occurs within weeks, confirming the formation of structurally mature myelin segments. Direct integration of MyeliMAP with HD-MEA enabled: (1) longitudinal assessment of OPC-mediated neuronal support to form functionally stabilized network; and (2) neuronal maturation in terms of myelin-dependent conduction dynamics alongside classical morphological readouts. The MyeliMAP-HD-MEA platform revealed progressive increase in neuron network maturation and axon conduction velocity, indicative of myelin-dependent signal propagation. These findings validate the MyeliMAP system as a physiologically relevant platform that captures both the morphological and electrophysiological signatures of human myelination *in vitro*.

## Materials and Methods

### hPSC culture

Two hPSC lines (biosafety documents: AMV/22042013 and SBB219.2012/1000), Sigma-0028-iSOX10 iPSCs (iPSC EPITHELIAL-1, Sigma, female, RRID: CVCL_EE38) and hDFa90/1.2-iNGN2 iPSCs were used in this study. hPSCs were maintained on hESC qualified Matrigel (Corning) coated 6-well plates in E8 flex medium (E8 basal medium with E8 supplement flex and 1% penicillin/streptomycin (Gibco)). hPSCs were passaged twice a week with 0.5 mM EDTA. The cells were regularly tested for mycoplasma contamination using the MycoAlert Mycoplasma Detection Kit (Lonza).

### iNGN2 neuron differentiation

iNGN2 neurons were generated by Novartis by overexpression of the transcription factor NGN2 in hDFa90/1.2 iPSC line, as described by *Russel et al*.^12^. Briefly, hiPSC line hDFa90/1.2-iNGN2 hPSCs were cultured in mTeSR, dissociated and plated at a density of 400,000 cells/cm2 on Matrigel-coated 6-well plates. Doxycycline 1 μg/ml in proliferation medium (Table S1) was added for 3 days to induce neuronal differentiation. The resulting neuron progenitors were cryopreserved in aliquots containing 2-3× 10⁶ cells in fetal bovine serum supplemented with 5% DMSO. These cells were kindly provided by Novartis Institutes for Biomedical Research, Switzerland.

### iSOX10 OPCs differentiation

iSOX10 OPCs were generated by overexpression of the transcription factor SOX10 in Sigma-iPSC0028 as described in *Garcia et al*.^17^. Briefly, Sigma-iSOX10-iPSC0028 were cultured in E8 flex medium, dissociated with Accutase and plated at a density of 25,000 cells/cm2 on a Matrigel-coated 6-well plate. After 1 day, oligodendrocyte induction medium (OIM, Table S1) was added for the following 6 days. On day 7, OIM containing 1 μM SAG was added and the medium was refreshed daily until day 11. Cells were then dissociated with Accutase and replated at 50,000-75,000 cells/cm2 in the Basal Medium (Table S1) on PLO-laminin coated 12-well plate to direct them towards the oligodendrocyte fate. The medium was changed every day until day 23. On day 24, the cells were dissociated with Accutase and cryopreserved in the same medium with 15% DMSO for further experimental use.

### Preparation of MyeliMAP on glass-bottom dish/HD-MEA chip

The PDMS networks were designed in AutoCAD (Autodesk) and fabricated by WunderliChips GmbH (Zurich, Switzerland). As illustrated in Figure 1, each design features a wide central axonal channel flanked by two peripheral seeding wells, separated by arrays of seven microchannels per side. The central axonal channel is 150 µm wide and 1.96 mm long, while the peripheral seeding wells measure 300 µm in width and 250 µm in length. The microchannels are 4 µm wide, tapering to 1.5 µm at their narrowest point to restrict the translocation of neuronal somata. Each peripheral seeding well terminates in a 30 µm-wide axon-collecting channel, allowing axons to converge. Following fabrication, PDMS networks were manually cut from the wafer, immersed in isopropanol for 15 minutes, rinsed three times with sterile water, and air-dried at room temperature (RT).

**Figure 1:**
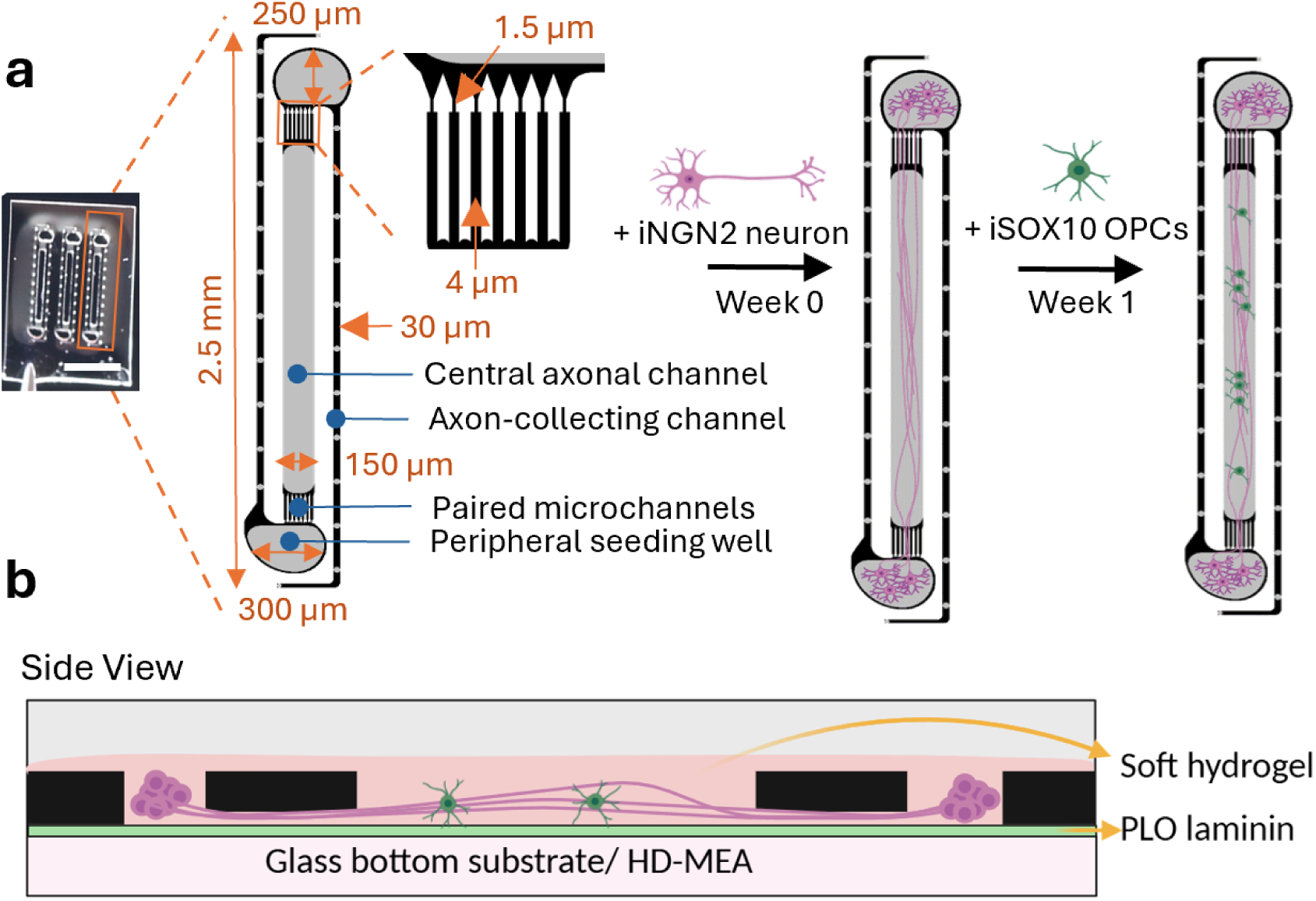
Schematic of MyeliMAP, an engineered functional myelinated networks-on-chip platform. **a** PDMS microstructure containing the MyeliMAP network geometry. Each unit consists of two peripheral seeding wells at opposite ends (250 × 300 µm), each connected to a central axonal channel (1960 × 150 µm) through paired microchannels on both the ends, having seven separate microchannels—1.5 µm wide at the corners and 4 µm wide at the center. The sequential axon-selection design of the microchannels ensures somata exclusion. The peripheral seeding wells are flanked by two 30 µm-wide axon-collecting channels. The grey regions of MyeliMAP are open on top to allow cell seeding and extracellular matrix (ECM) filling, while the underside (black region) contains electrodes for electrophysiological recordings. iNGN2 neuron spheroids are seeded in the peripheral seeding wells and allowed to extend axons into both the central axonal channel and axon-collecting channels for one week, after which oligodendrocyte progenitor cells (OPCs) spheroids are seeded into the central axonal channel. **b** Side view of MyeliMAP integrated onto either a glass substrate or a high-density microelectrode array (HD-MEA). Neuronal spheroids extend axons into the central axonal channel within a low concentrated ECM, where they establish interactions with oligodendrocytes. Scale bar: 2mm (a).

Before applying the microstructure on a glass-bottom dish (KIT-3522 T, WillCo Wells)/HD-MEA chip (MaxOne+ Chips, Pt-electrodes, Maxwell Biosystems), the glass/HD MEA chip surface was coated with poly-L-ornithine for 1 hr and laminin overnight. Then, microstructures were added to either the glass bottom dish for immunofluorescence imaging or the HD-MEA chip for electrophysiology and immunofluorescence imaging.

### Loading of Matrigel based extracellular matrix (ECM) in MyeliMAP

Human Matrigel (stock concentration-8.7mg/ml) was diluted 1:4 in proliferation medium and supplemented with iMatrix laminin (final concentration ∼70 μg/ml). The mixture was carefully applied onto the PDMS microstructure positioned on either a glass-bottom dish or an HD-MEA chip using a 10 μl pipette. To promote even distribution, the gel was gently agitated to allow migration into the microchannels, and excess material was removed with the same pipette tip to minimize cell overgrowth on the surface of the MyeliMAP. The MyeliMAP with gel was then incubated at 37 °C for 10 minutes (min) and phosphate buffer saline (PBS) was added to the dish/chip until spheroid seeding.

### iNGN2 neurons and iSOX10 OPC seeding as spheroids in MyeliMAP

iNGN2 neurons were thawed at 37 °C, transferred dropwise into 4 mL of pre-warmed neuronal maintenance medium (NMM, Table S1), followed by centrifugation at 300 relative centrifugal force (RCF) for 5 min. Cells were counted and 300,000 cells (400 cells/spheroid) were added to spheroid forming plate (spherical plate 5D®, Kugelmeiers®, SP5D-24W), pre-filled with 1 mL of warm NMM supplemented with RevitaCell. Next, the spheroid plate with the iNGN2 neurons was centrifuged at 50 RCF for 5 minutes and incubated overnight at 37 °C for spheroid formation.

iSOX10 OPCs were thawed at 37 °C, transferred dropwise into 4 mL of pre-warmed oligodendrocyte maturation medium (OMM, Table S1) medium and centrifuged at 300RCF for 5 min. 100,000 cells/cm2 iSOX10 OPCs were plated on PLO-laminin coated 6-well plate in OMM supplemented with 5 μg/ml doxycycline. Medium exchanged with fresh OMM with doxycycline was done for a week, iSOX10 OPCs were detached using accutase and spheroids were formed as described above for iNGN2 neurons.

The PDMS microstructure containing the Matrigel based ECM was first incubated with PBS and then desiccated until all bubbles were eliminated. Following this, PBS was carefully removed, and NMM supplemented with RevitaCell was added onto the prepared glass-bottom dish or HD-MEA chip. Spheroids of iNGN2 neurons were seeded in the two peripheral seeding wells of each of the three networks (one spheroid/well) using a 10μL pipette and the glass bottom dish/HD-MEA chip with iNGN2 neuron spheroids was carefully transferred to incubator. iNGN2 neurons were first cultured in NMM for one week, after which iSOX10 OPC spheroids were introduced into the central channel of each of the three networks (three spheroids per channel). The following day, the medium was replaced with OMM supplemented with 3 µg/ml doxycycline. One week after iSOX10 OPC addition, the medium was again replaced with OMM. Thereafter, half of the medium was exchanged twice weekly for an additional 6-8 weeks.

### Immunofluorescence

The glass-bottom dishes/HD-MEA chips were fixed at different timepoints (weeks 2-8) with 4% phosphate buffered paraformaldehyde for 10 min at RT. After fixation, samples were washed with PBS, blocked and permeabilized using 0.5% Triton X-100 in PBS (PBST) supplemented with 5% normal goat serum (Dako) for 1 hour at RT. Samples were incubated with primary antibodies (Table S2) diluted in PBST overnight at 4 °C in humidified chambers, washed with PBS three times with 5 min interval, and secondary antibodies (species matched) diluted in Dako Real Antibody diluent were added for 1 hour at RT. Samples were washed with 0.01% PBST, three times with 10 min intervals, and nuclei were stained with Hoechst 33342 (Sigma, 1:1000 dilution). The immunostained samples were imaged using a confocal laser scanning microscope (CLSM) (Fluoview 3000, Olympus).

### Transmission Electron Microscopy (TEM)

Glass-bottom dishes with cells in microstructures at week 6 were fixed in 2.5% glutaraldehyde (Electron Microscopy Services) in 0.1 M sodium cacodylate buffer (Electron Microscopy Services), pH 7.6 for 45 min at RT. Samples were washed thrice with 0.1M sodium cacodylate buffer and 1% osmium tetra-oxide (Electron Microscopy Services) with 1.5% potassium ferrocyanide (Sigma) diluted in 0.1 M sodium cacodylate buffer was added. Samples were incubated for an hour at 4 °C on a shaker. The samples were washed with Milli-Q water six times at 2 min intervals, followed by overnight incubation in 2% Uranyl acetate (Electron Microscopy Services) diluted in Milli-Q water at 4°C. The next day, samples were washed five times for 7 min with Milli-Q water and dehydration was done by increasing concentrations of ethanol (30%, 50%, 70%, 90%, 100%) on ice, 10 min each. Epon resin/ethanol mixtures was added to the samples, before overnight infiltration with pure epon resin (Agar Scientific). The following day, a beam capsule with fresh epon was added carefully on the top of microstructures and samples were incubated at 60 °C for 48 hours. Next, the glass was removed from the samples with quick exposure to liquid nitrogen, 80nm thick longitudinal and cross-sections were made using a diamond knife (Diatome Ltd.) and Leica UC7 ultramicrotome (Leica Microsystems),and placed on Formvar/carbon coated TEM grids (Quantifoil). Sections were stained with 2% aqueous uranyl acetate for 15 minutes and Reynold’s lead citrate for 5 minutes. Micrographs of the stained sections were imaged using the Talos L120C TEM (Thermo Fisher Scientific) equipped with a camera BM-Ceta, operating at 120 kV acceleration voltage in brightfield mode camera.

### Electrophysiology

Electrophysiology recordings were performed as described in^18,19^. Briefly, the MaxOne MEA chip (Maxwell Biosystems) was placed on the recording unit in an incubator with 35°C air temperature, 90% humidity and 5% CO2 concentration. Recordings were only initiated five minutes after placement of the HD-MEA chip placement in incubator, ensuring that any spontaneous activity disrupted by movement or abrupt changes in CO2 levels and temperature due to handling could return to baseline. Each individual network per PDMS microstructure were identified and selected utilizing the impedance map method, extensively described in ^18–20^. This method allows electrode identification covered by PDMS microstructure as opposed to electrodes exposed to the conductive medium solution via a threshold applied on the measured impedance. Electrodes of interest underlying medium-exposed network regions for spontaneous recordings and electrodes under the 7 lateral microchannels were manually selected for stimulation induced activity measurements) were routed, and network activity was tracked by recording the voltage on the routed electrodes.

#### Spontaneous recordings and analysis

Voltage recordings were acquired at 20 kHz sampling frequency for 2 min weekly from electrodes of interest, with a resolution of 10 bits and a recording range of approximately +/- 3.2 mV, which results in a least significant bit (LSB) corresponding to 6.3 µV. Using custom software based on the MaxWell Python API^20^, the raw traces were recorded and stored as HDF5-files. Raw hdf5 files were processed with a custom Python pipeline. Spikes were detected and sorted with the SpikeInterface framework^21^ using the SpyKING CIRCUS v2 sorter^22^ to calculate single neuron parameters. Detection thresholds were set to negative peaks exceeding 5 × signal standard deviation, with a minimum interspike interval (ISI) of 3 milliseconds (ms). Identified neuronal units were discarded if their signal-to-noise ratio was < 5 or their firing rate < 0.1 Hz. For each retained neuron, firing rate and burst rate were calculated. Bursts were identified using an ISI-based method: a burst consisted of ≥ 4 consecutive spikes within ≤ 100 ms and was terminated when this condition was no longer met. To quantify network-level activity, network bursts were quantified across all neurons by threshold-based peak detection in the population firing rate. The population firing rate was obtained by binning detected spikes in 25 ms windows and smoothing with a Gaussian kernel. Network bursts were defined as peaks in the population firing rate exceeding the baseline activity by 7 standard deviations for ≥ 50 ms, involving at least 30 % of identified neurons. Baseline activity was estimated as the median of the lower 50% of the smoothed population firing rate. Network bursts ended when activity dropped below 1 standard deviation above the baseline activity for ≥ 50 ms. To avoid false positives in low-activity cultures, the minimum network burst onset threshold was set to 6 Hz.

#### Stimulation recordings and analysis

For stimulation-induced activity, a repetitive stimulation pattern was applied to a microstructure, via a custom Python script. The script makes use of the MaxWell Python API to send commands to the system hub. A biphasic pulse with a leading cathodic phase and 400 µs pulse width was used with a stimulation amplitude of 1000 mV (2000 mV peak-to-peak). The amplitudes were rounded to the closest value available on the 10-bit DAC with approximately 3.2 mV step size. The stimulation (500 pulses, 4 Hz, 1000 mV voltage amplitude), was applied first to the 7 lateral microchannels on one side of the central axonal channel of each network, followed by the contra-lateral set of 7 microchannels per network. Electrophysiology measurements were always performed the day after last medium exchange. In the longitudinal experiment, 6-15 networks per condition (distributed across 3 networks per chip) were recorded for their spontaneous activity and stimulation-based activity every week from week 1 to 9.

Analysis of evoked responses was performed according to the analysis pipeline developed in^19^. Briefly, every single lateral microchannel was visually identified on the impedance scan and one electrode per microchannel was manually selected via a custom-made GUI and saved in a separate file (located in the middle of the microchannel itself). The post-stimulus raster plots were generated for each electrode (one per microchannel) to visually inspect the response of axons in each microchannel to the various stimulation protocols. As the stimulation signal saturates the amplifiers (capped at 3.2 mV) independently from the stimulation amplitude, an artifact amplitude threshold of 3 mV was established to detect the stimulation pulse times on the stimulation electrode and was used a control for checking that stimulation was adequately delivered. In case of defective stimulation with missing artifact for all recording timepoints, the corresponding networks were excluded from the analysis (2 coculture network). For each selected electrode per microchannel, spikes were detected within 0.5 ms and 30 ms from the stimulation artifact. This time window was sufficiently long for the action potentials to travel from the stimulation electrode to the microchannels. Raw data were processed using a band-pass filter (4th order acausal Butterworth filter, 300-3500Hz). The baseline noise of the signal was characterized using the median absolute deviation for each electrode. Action potential event times were extracted from the voltage trace by identifying negative signal peaks below a threshold of 5 times the baseline noise. Successive events within 0.75 ms were discarded to avoid multiple detection of the same spike. Spike amplitudes were defined as the absolute value of the negative amplitude of the detected peak. As described in more details in^19^, each induced response upon nerve stimulation was extracted via a clustering algorithm; and metrics such as number of responses, conduction speed. The conduction speed was calculated from the latency, defined as the time delay between the stimulus and the induced response at the recording site, at the beginning of the stimulation. The conduction speed was then obtained by dividing the distance by the latency. The distance represents the hypothetical path taken by the axon from the recording site to the stimulation site and was approximated to be 1960µm for all data points, based on the theoretical measurement of shortest possible path and longest possible path.

### Statistical analyzes

Each MyeliMAP microstructure plated as independent experiment represent a biological replicate (N) having three networks within that microstructure as technical replicates (n). For spheroid morphological analysis, number of independent spheroid batches used represents biological replicates and number of spheroids in each batch represents technical replicates. The number of these biological (N) and technical (n) replicates is mentioned in the respective figure legends Statistical analyzes were performed using Prism v.8.2.1 (GraphPad). Tests used include two-way ANOVA with Turkey’s multiple comparisons test, one-way ANOVA with Turkey’s multiple comparisons test, and are specified in the figure legends. Differences were considered statistically significant for two-sided p-values < 0.05 (*p < 0.05; **p < 0.01; ***p < 0.001; ****p < 0.0001).

## Results

### 1. MyeliMAP’s microengineered architecture facilitates *in vivo*-like neuron myelination

We developed MyeliMAP, a microphysiological platform designed to model human myelination *in vitro* by combining monocultures or cocultures of human-derived iNGN2 neurons with or without iSOX10 oligodendrocyte precursor cells (OPCs) within a microengineered PDMS-based microstructure. Each microstructure contained three identical networks, allowing for medium-throughput experimentation through multiplexing (Figure 1a). The microstructure design provided precise spatial control over neuronal organization and neuron-glia interactions, closely mimicking the architecture of human white matter tracts^23^. Within each network, two peripheral seeding wells (250 × 300 µm) received iNGN2 neuronal spheroids, from which neurites extended along two distinct paths: (1) a 30 µm-wide axon-collecting channel that funneled fibers into a dense, aligned axonal bundle, and (2) a set of two paired microchannels containing seven soma-excluding microchannels (widening from 1.5 µm to 4 µm) that prevented somatic migration while permitting distal neurite projection. These microchannels converged into a central neurite/axonal channel (1960 × 150 µm), where neurites/axons from opposite wells met and formed aligned tracts. One week after neuronal seeding and neurite extension, iSOX10 OPCs were introduced into the 150μM central channel, where they codifferentiated alongside the preformed axonal bundles to generate compact myelin sheaths. The defined PDMS network designs provided micron level spatial precision, overcoming the poor compartmentalization and variability typical of conventional coculture systems. MyeliMAP could be integrated with either glass-bottom plates or high-density microelectrode array (HD-MEA) substrates (Figure 1b), enabling parallel structural (immunofluorescence) and functional (electrophysiological) analyses. The system supported long-term cocultures (> 70 days), allowing coordinated neuronal and oligodendrocyte maturation and sustained *in vitro* myelination. The PDMS-guided configuration imposed an *in vivo*-like separation between neuronal somata and axonal tracts while maintaining full experimental accessibility. Delayed OPC introduction ensured neurite outgrowth prior to myelination, thereby enabling temporally and spatially regulated axon-glia interactions that closely recapitulate the dynamics of myelination *in vivo*^24^. Collectively, these design features establish MyeliMAP as a powerful human-based platform for modeling activity-dependent myelination, bridging structural precision with functional relevance.

### 2. Optimized Matrigel based ECM supports neuron and oligodendrocyte codifferentiation and maturation

3D culture systems more effectively support *in vitro* myelination than traditional two-dimensional (2D) cultures, as they better reproduce the spatial architecture and intercellular interactions of the central nervous system^25–28^. Within these 3D environments, ECM components such as Matrigel and laminin provide essential biochemical cues that mimic a myelin-supportive niche, thereby promoting neurite outgrowth, OPC differentiation, and axon ensheathment^29–31^. In MyeliMAP, we optimized a 3D Matrigel based ECM formulation by diluting Matrigel in culture medium and supplementing with laminin to balance neuronal survival with oligodendrocyte support. To standardize the placement of neuronal spheroids inside the peripheral seeding wells, 400-cell neuronal spheroids (∼170 μm diameter) and iSOX10 OPC spheroids of comparable size were used (Figure S1a–b). Neurons survived in both 1:1 and 1:4 Matrigel dilutions; however, while 1:1 Matrigel favored cell clustering, 1:4 Matrigel promoted extensive neuronal arborization and enhanced OPC survival. Diluted Matrigel based ECM formulation thus provided a more favorable environment for oligodendrocyte viability and growth, enabling reliable coculture with neurons for myelination studies (Figure S1c).

### 3. MyeliMAP enables neuron myelination

We employed the MyeliMAP configuration with the optimized 1:4 Matrigel based ECM formulation to sequentially coculture iNGN2 neurons and iSOX10 OPCs in the peripheral seeding wells and central axonal channel, respectively. iNGN2 neuron spheroids placed in the peripheral seeding wells extended cell extensions through the microchannels into the axon-collecting channel within one week. iSOX10 OPC spheroids introduced into the central axonal channel migrated outward within a day and elaborated processes in close proximity to neuronal axons within a week (Figure 2a). By five weeks, we observed dense oligodendrocyte arborization (MBP+) colocalizing with neuronal axons (NF+) across the central axonal channel, consistent with progressive oligodendrocyte maturation and axonal ensheathment (Figure 2b,c). This was further supported by the colocalization of nodes of Ranvier (CASPR+ paranodes) at contact sites between axons (NF+) and oligodendrocyte projections (MBP+) (Figure 2d). These findings highlight the unique capacity of MyeliMAP to integrate human-derived neural cells within a physiologically relevant segregated microenvironment, enabling the spatially relevant study of myelination.

**Figure 2:**
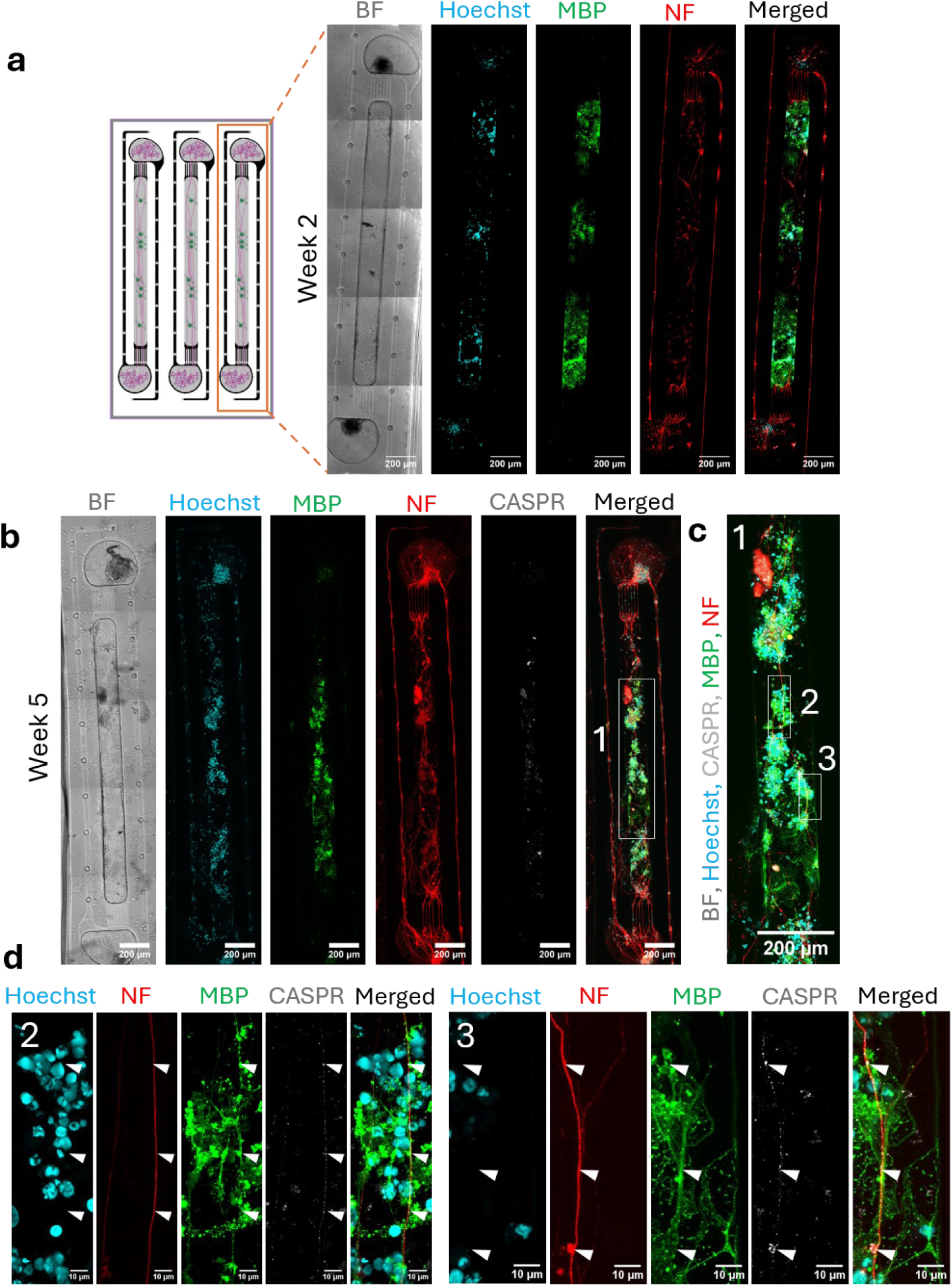
MyeliMAP supports co-maturation of neurons and oligodendrocytes to enable physical interactions for myelin formation. **a** PDMS device containing three MyeliMAP microstructures. Confocal images obtained after two weeks in culture show neuronal spheroids (NF+) seeded in peripheral compartments projecting axons into the central lane, where they interact with oligodendrocytes (MBP+). Brightfield (BF) images and nuclear staining (Hoechst) are included for reference. **b** After 5 weeks of culture, immunofluorescence imaging reveals enlarged oligodendrocyte structures (MBP+) and alignment of neuronal (NF+) axons with oligodendrocyte (MBP+) processes, accompanied by expression of the node of Ranvier marker CASPR+. BF and Hoechst images are shown for context. **c,d** Higher magnification of panel b highlights aligned neuron-oligodendrocyte structures (c, region 1 from panel b) and colocalization of CASPR+ nodes of Ranvier on these structures (indicated with white arrowheads) (d, region 2 and 3 from panel c). Scale bar: 200 μm (a,b,c) and 10 μm (c). Representative immunostaining images for N=3 (a-d).

To further validate myelination within MyeliMAP, we performed ultrastructural analysis using TEM on six-week cocultures (Figure 3a). TEM imaging revealed axons at distinct stages of myelin development, ranging from initial, loosely wrapped sheaths (Figure 3a-b) to mature, multilayered compact myelin (Figure 3c-d). The observation of progressive wrapping and compaction around human axons confirmed that oligodendrocytes in MyeliMAP undergo the full spectrum of myelination events *in vitro*. These ultrastructural features provided direct evidence for functional myelination, complementing the immunofluorescence findings of MBP+ oligodendrocyte processes colocalizing with NF+ axons and CASPR+ paranodes. Together, these results demonstrate that MyeliMAP is a robust, long-term human *in vitro* platform to study oligodendrocyte maturation, axonal ensheathment, and myelin sheath stabilization under defined microphysiological conditions.

**Figure 3:**
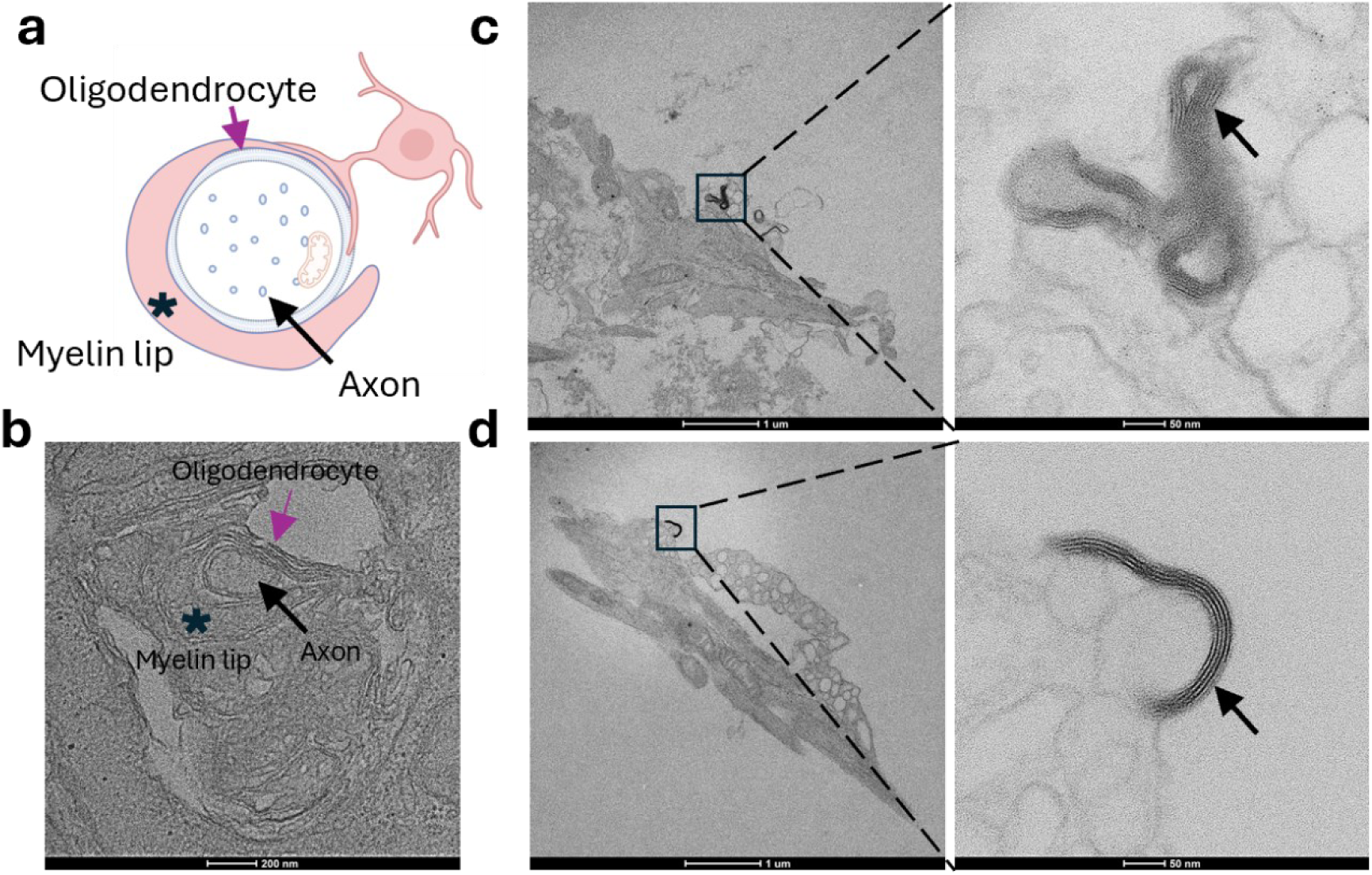
Ultrastructural validation of neuron myelination in MyeliMAP with neuron-oligodendrocyte coculture. **a** Schematic illustration of neuron-oligodendrocyte interactions within MyeliMAP showing initial loose myelin wrapping at week 6 in culture. **b** Transmission electron microscopy (TEM) imaging confirmed loose myelin wrapping corresponding to the schematic in panel a. Black arrow: axon; Purple arrow: oligodendrocyte; Asterisk: myelin lip (a,b). **c,d** TEM images also demonstrated the presence of compact myelin sheaths (indicated with black arrows) at week 6 in culture. Scale bars: 200 nm (b), 1 µm and 50 nm in zoomed panels (c,d). Representative images from N = 2 independent experiments on MyeliMAP with neuron-oligodendrocyte coculture with n=6 networks.

### 4. Oligodendrocytes enhance neuron network maturation in MyeliMAP

To assess the functional impact of oligodendrocytes on neuronal maturation, we compared iNGN2 neuron monocultures with iNGN2 neuron-iSOX10 OPC cocultures using the integrated MyeliMAP-HD-MEA platform (Figure 4a). Spontaneous electrophysiological activity was recorded weekly for nine weeks, and both single neuron parameters (firing rate and bursting rate) and network-level parameters (network bursting rate, network burst spike rate and percentage of active neurons per network burst) of neuron monoculture and neuron-oligodendrocyte coculture in MyeliMAP-HD-MEA were analyzed. During the early culture period (weeks 1-4), neuron monocultures exhibited higher single cell firing and bursting rates relative to cocultures (Figure 4b-c). However, neuronal activity in mixed cultures progressively increased and, by weeks 8-9, exceeded that of monocultures, with significantly higher firing and bursting rates. At the network level, oligodendrocyte cocultures showed increasing network bursting and network-burst spike rates from weeks 5-9, whereas monocultures began higher but declined over time (Figure 4d-e). Likewise, the fraction of neurons recruited per burst rose steadily in coculture but plateaued in monocultures, indicating oligodendrocyte-dependent neuronal network recruitment (Figure 4f). Week 9 MyeliMAP-HD-MEA platform containing neuron monocultures and neuron-oligodendrocyte cocultures was stained for neuronal (NF+) and oligodendrocyte (MBP+) markers (Figure 4g,h). This confirmed the presence of oligodendrocytes and their aligned morphology along neuronal axons in the coculture condition, demonstrating active neuron myelination on MyeliMAP-HD-MEA platform (Figure 4g,h and Figure S2-3). Collectively, these findings indicate that oligodendrocytes progressively stabilize and enhance neuronal firing properties and promote more robust and sustained network activity over prolonged culture periods.

**Figure 4:**
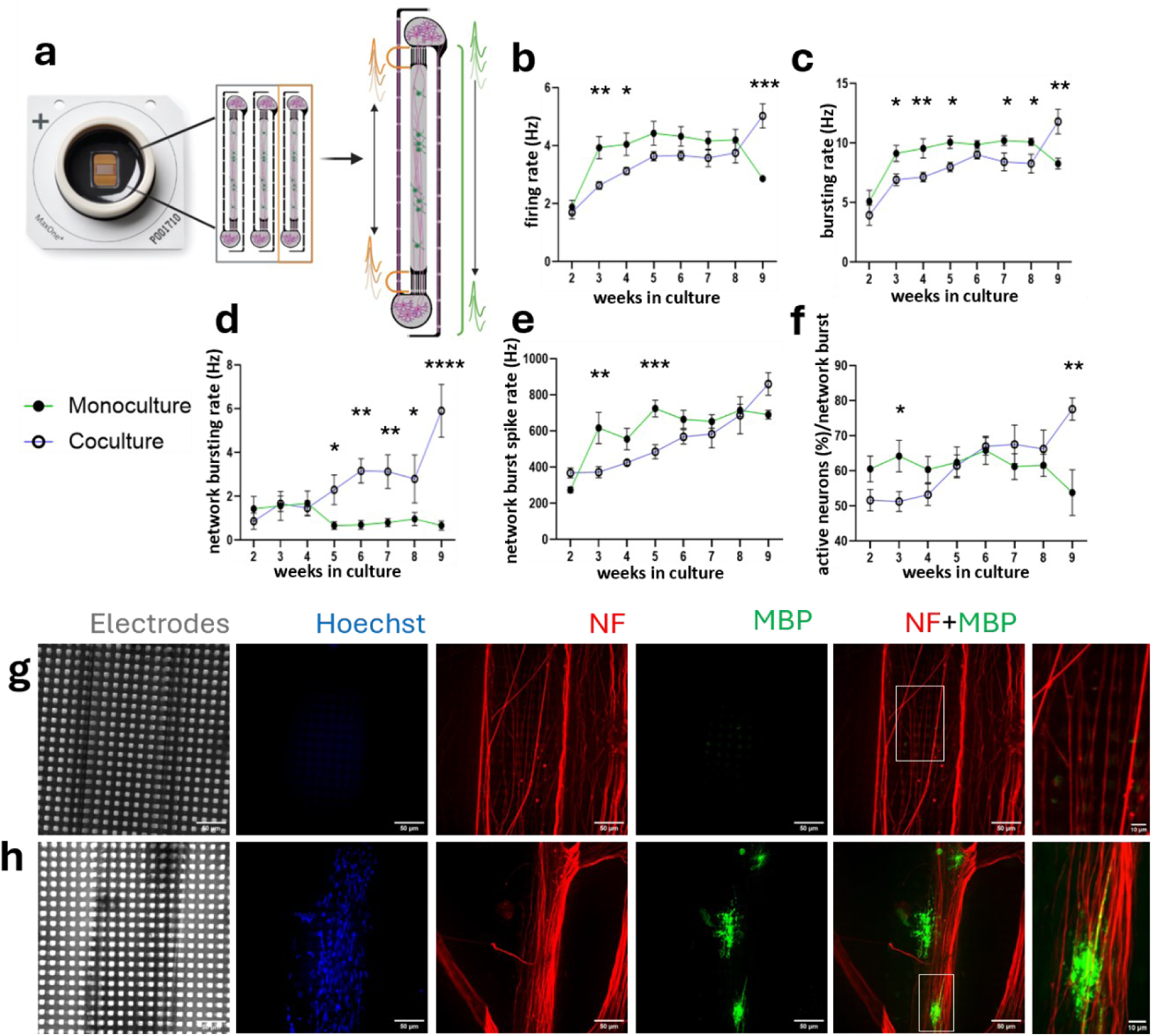
Oligodendrocytes reduces hyperexcitability and improves network synchronization of neurons. **a** Schematic illustration of MyeliMAP with neuron-oligodendrocyte coculture on a high density-microelectrode array (HD-MEA). The grey regions are open-on-top exposed with culture medium, and the black regions contain electrodes used for recordings and stimulations. Action potential conduction is bidirectional in the central axonal channel (indicated with orange color) and unidirectional in the axon-collecting channels (green color). **b-c** Single neuron parameters (firing rate, bursting rate) of neuron monoculture (green line) and neuron-oligodendrocyte coculture (blue line) reveals early hyperexcitability in monocultures (weeks 1-4), whereas cocultures progressively increased activity and surpassed monocultures by weeks 8-9. **d-f** Network parameters (network bursting rate, network burst spike rate and percentage of active neurons per network burst) of neuron monoculture (green line) and neuron-oligodendrocyte coculture (blue line) shows that cocultures develops stronger network bursting, higher spike rates per burst, and greater neuronal recruitment in network burst from weeks 5-9, while monoculture activity declined. **g-h** Immunofluorescence imaging reveals neuron (NF+)-oligodendrocyte (MBP+) alignment in coculture compared to neuron monoculture in week 9 MyeliMAP-HD-MEA platforms. Brightfield and Hoechst images are shown for context. Scale bar: 50 μm and 10 μm in zoomed panels (g,h). Data shown as mean ± SEM. Statistical analysis were performed by Two-way ANOVA with Turkey Multiple Comparison Test (b-f). N=5 independent experiments (HD-MEA chips) with n=15 networks (week 2-8); N=2 independent experiments (HD-MEA chips) with n=6 networks (week 9) (b-f). . *p< 0.05, **p< 0.01, ***p< 0.001, ****p0.0001.

### 5. Neuronal myelination enhances action potential conduction in MyeliMAP

To quantitatively assess myelination-dependent functional changes in MyeliMAP, we compared action potential conduction in iNGN2 neuron monoculture and iNGN2 neuron-iSOX10 OPC coculture using the integrated MyeliMAP-HD-MEA platform. Stimulation was applied once a week during the recordings from 2-8 weeks, to each paired microchannel set featuring seven parallel microchannels laterally positioned on either side of the central axonal channel of each microstructure (500x pulses, 4 Hz, +/-1000 mV voltage amplitude). Evoked responses were recorded at the contra-lateral paired microchannel set at each of the seven microchannel and vice versa (Figure 5a). For each condition, both the number of responses (reflecting recruited axons present near the recording electrodes) and the latency of signal propagation were measured (Figure 5b). As the distance between the paired microchannels was defined by the fixed microchannel geometry, conduction velocities were calculated for weeks 2-8. Neuron monocultures consistently showed a higher number of responses compared to cocultures, likely reflecting differences in axon excitability or axonal density (Figure 5c). Importantly, latencies were initially similar between conditions but progressively decreased in cocultures, resulting in significantly faster conduction velocities from week 6 onward; the time point at which compact myelin was detected. In contrast, latency and conduction velocity in monocultures remained unchanged (Figure 5d-e). These results demonstrate that addition of oligodendrocytes in MyeliMAP not only promotes structural myelination but also drives functional improvements in axonal conduction associated with neuronal maturation.

**Figure 5:**
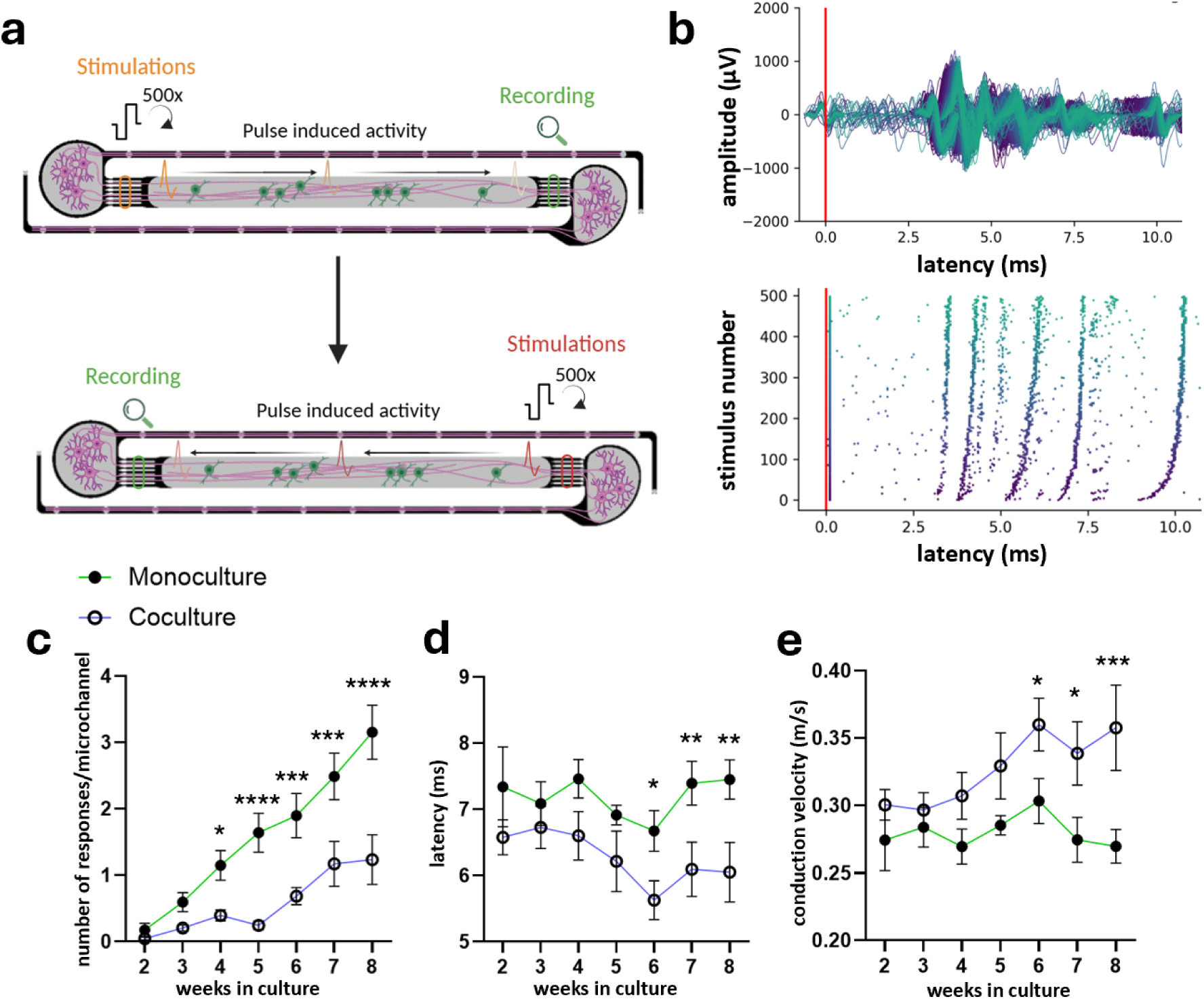
Myelination in MyeliMAP enhances action potential conduction. **a** Schematic of the stimulation protocol: sequential pulses (500x, 4 Hz) were delivered through electrodes beneath one end of a paired microchannel, while responses were recorded from electrodes at the opposite end and vice versa. **b** Post-stimulus raster plot showing representative responses in a single microchannel, illustrating stimulus-induced activity amplitude and latency over repeated pulses. **c** Quantification of the number of responses per microchannel in neuron monocultures (green) and neuron-oligodendrocyte cocultures (blue). **d-e** Latency measured across conditions (d) are used to calculate conduction velocity (e) based on the fixed central axonal channel length from weeks 2-8. Data shown as mean ± SEM. Statistical analysis was performed by Two-way ANOVA with Turkey Multiple Comparison Test (c-e). N=5 independent experiments (HD-MEA chips) with n=15 networks (b-e). *p< 0.05, **p< 0.01, ***p< 0.001, ****p0.0001.

## Discussion

Human-based myelination models, such as organoid systems, are essential for capturing species-specific features of oligodendrocyte development and pathology that rodent models cannot fully recapitulate. However, only a few human *in vitro* myelination platforms currently exist, and those that are available typically require more than 10 weeks to achieve neuronal myelination, while still lacking functional readouts to study neuron-oligodendrocyte interactions^32–35^. Further, myelination is a complex process that depends on oligodendrocytes extending and wrapping their membranes around neuronal axons, effective axonal isolation and a physiologically relevant ECM are crucial^36^—features that conventional systems often do not fully provide. Microfluidic technologies have long been used to separate axons from their cell bodies to enable controlled axon–glia interactions. A landmark example by Kerman et al., 2015 demonstrated *in vitro* myelination using mouse embryonic neurons and oligodendrocytes, providing valuable insight into axon ensheathment and myelin compaction^37,38^. However, this model relied on rodent cells, used relatively large microchannels (2–5 μm) that allowed multiple axons to pass simultaneously, and lacked functional readouts—factors that limit its translational relevance.

To address these limitations, we developed MyeliMAP, a platform that integrates PDMS-based, microstructure-guided architecture with HD-MEAs to create a myelination-permissive environment with simultaneous structural and functional readouts. MyeliMAP addresses these challenges by recreating an *in vivo*-like microenvironment with micron level spatial control of axonal networks and long-term culture compatibility in a myelination-permissive setting. To achieve this, we initially tested several previously described microstructure designs^18,39,40^ to guide axons and oligodendrocytes, aiming to ensure proper alignment and sufficient distance between stimulation and recording sites (>1 mm) to avoid stimulation artifacts and accurately measure conduction velocity. These established designs, however, proved inadequate: oligodendrocytes—being larger and less migratory than Schwann cells—struggled to move through narrow channels, and some layouts failed to preserve the required electrode spacing (data not shown)^18^. Through iterative optimization, we arrived at the configuration described here, featuring a central, widened lane that enables oligodendrocytes to exit spheroids efficiently without needing to migrate through the axon-collecting channel, while maintaining sufficient separation between stimulation and recording electrodes. This architecture supports both myelination and functional assessment and offers flexibility to introduce distinct neuronal populations into the two peripheral compartments. As a result, MyeliMAP provides a human-relevant platform for studying neuronal-subtype–specific myelination, neuron–glia interactions, and long-range connectivity between brain systems—such as modeling myelinated pathways like the optic nerve.

Beyond axonal isolation, the ECM plays a decisive role in modulating myelination. Previous studies have shown that oligodendrocytes are highly responsive to ECM components, where Matrigel and laminin based environments promote process extension, axonal ensheathment, and myelin formation^30,41–43^. Guided by these findings, we optimized a Matrigel based ECM supplemented with laminin to support OPC adhesion, differentiation, and maturation. We found that lower concentration of Matrigel based ECM supported better oligodendrocyte survival and extensions, in line with other studies showing benefits of softer matrices in axonal ensheathment and myelin formation^44,45^. Therefore, the presence of ECM proteins may provide essential cues to support initial OPC adhesion and growth. It is also likely that long-term culture may further alter the biomechanical properties of the gel through ongoing cell-matrix interactions, which was not yet assessed in this study. Nevertheless, the Matrigel based ECM formulation used here, in combination with the defined microarchitecture of MyeliMAP, supports robust oligodendrocyte growth and promotes codifferentiation of iNGN2 neurons and iSOX10 OPCs. To investigate neuron-oligodendrocyte interactions in MyeliMAP, we maintained cocultures of iNGN2 neurons and iSOX10 OPCs for six weeks, followed by immunofluorescence analysis. Mature, arborized MBP+ oligodendrocytes were observed in close alignment with NF+ neuronal extensions, indicating direct physical interactions between the two cell types. Notably, CASPR+ puncta were detected at points of contact, suggesting the formation of functional paranodes and compact myelin segments. However, this alignment was not uniform, and we observed only partial rather than complete myelination of the neuronal network. TEM further confirmed the presence of both loosely wrapped and compact myelin sheaths, as well as unmyelinated axons, indicating that not all neurons were myelinated at this stage (data not shown). This incomplete myelination may reflect the relatively immature state of iNGN2-derived neurons, which likely require additional somatic and axonal maturation before becoming fully permissive for oligodendrocyte wrapping^46^. It may also result from the limited number of mature oligodendrocytes present around neurites in the central axonal channel, as supported by the high proportion of immature oligodendroglial cells (MBP negative) observed in Figure 4h and S3 and by previous studies reporting broad heterogeneity in oligodendroglial maturation profiles in neuron-glia coculture systems^6,47^. Further studies, where the co-culture time is extended, or where maturation of neurons and/or oligodendrocytes is further enhanced by modifying culture media might overcome these shortcomings. The co-maturation of neurons and oligodendrocytes in MyeliMAP is also strongly influenced by the ECM composition. Although the Matrigel based ECM supported iSOX10 OPC survival, growth, and structural interactions with neurons, it may still lack specific signaling cues such as myelin-supporting proteins and lipids; required for complete and uniform neuronal myelination. Together, these results demonstrate that MyeliMAP supports a heterogeneous culture environment containing both myelinated and unmyelinated axons, effectively recapitulating the progressive and regionally variable nature of cortical myelination observed *in vivo*^48^.

In addition to providing defined structural geometry and a physiologically relevant ECM for permitting *in vitro* neuron myelination, a key advantage of MyeliMAP is its integration with HD-MEA technology, which enables quantitative evaluation of the functional crosstalk between iNGN2 neurons and iSOX10 OPCs. Using iNGN2 neuron monocultures and iNGN2 neuron-iSOX10 OPC cocultures, we demonstrated that iSOX10 OPCs modulate neuronal electrophysiology at both the single-neuron and network levels. At the single-cell level, spontaneous firing and bursting rates were initially higher in monocultures, reflecting greater excitability. In contrast, activity in cocultures progressively increased over time and eventually matched or exceeded that of monocultures, suggesting differential neuronal maturation in the presence of iSOX10 OPCs. At the network level, parameters including network bursting rate, network burst spike rate, and the percentage of active neurons within network bursts all demonstrated that oligodendrocytes enhance the synchronization and stability of neuronal activity, likely through progressive myelination. In line with these findings, Mazuir et al. showed that OPCs reduced excitability of GABAergic neurons but improved network coordination and increased the frequency of synaptic events, supporting the idea that OPCs do not simply suppress activity but refine it to produce more efficient network function^49^. Similarly, other recent studies have demonstrated that OPCs in close proximity to neuronal axons can attenuate neuronal firing rates while contributing to long-term stabilization of network dynamics, highlighting their dual role in dampening excessive excitability while fostering coordinated activity patterns^50^. Collectively, these reports underscore the emerging view of OPCs as active regulators of circuit physiology rather than passive precursors of myelinating oligodendrocytes. Within this context, MyeliMAP provides a powerful human-based model to dissect how OPCs and oligodendrocytes shape both the maturation and functional integration of neuronal networks.

Finally, the MyeliMAP platform integrated with HD-MEA technology was used to quantitatively assess functional improvements in neuronal action potential conduction mediated by oligodendrocytes. Comparing iNGN2 neuron monocultures with iNGN2 neuron-iSOX10 OPC cocultures, we observed a greater number of responsive signal bands in neuron-only cultures, likely reflecting differences in axonal arrangement. This may be due to the physical presence of iSOX10 OPCs in the coculture, which can partially obstruct axonal trajectories or alter their spatial distribution. We next calculated conduction velocity based on latency measurements between stimulation and recording electrodes across the defined channel geometry. Strikingly, conduction speed was significantly higher in iNGN2 neuron-iSOX10 OPC combined cultures compared to monocultures after week 6, coinciding with the appearance of compact myelin. This finding demonstrates that oligodendrocyte-mediated myelination in MyeliMAP translates into measurable improvements in axonal signal propagation. However, the absolute conduction velocities were lower than those reported in other comparable *in vitro* systems^18,51^. We attribute this to three main factors: (i) the open design of the central axonal channel, which allows axons to grow in non-linear trajectories rather than in tightly confined paths, thereby reducing conduction efficiency between stimulating and recording electrodes; (ii) the residual Matrigel based ECM layer on the microstructure surface, which can promote axonal growth outside the intended plane and thus create longer, less direct conduction routes; and (iii) incomplete myelination along axons, reflected by patchy or partially compacted myelin sheaths that reduce signal transmission efficiency. The first two limitations could be mitigated by refining the design to incorporate longer or more spatially restrictive microchannels^39^, applying plasma cleaning after Matrigel based ECM addition, or introducing anti-fouling surface coatings to prevent undesired cell overgrowth^51^—approaches proven effective in related microphysiological systems but not yet implemented in MyeliMAP. The third limitation of possible incomplete myelination can be addressed by performing TEM-based quantification of myelin architecture across multiple time points to track the transition from loose to compact myelin, determine the proportion of fully myelinated versus unmyelinated axons, and identify the optimal maturation window for functional measurements (as done in *Ahuja et al.*^6^). Together, these findings validate that MyeliMAP not only supports structural myelination but also enables quantitative assessment of myelin-dependent functional improvements in conduction velocity. This establishes MyeliMAP as one of the first human-based *in vitro* brain models to demonstrate iNGN2 neuron myelination featuring neuron-oligodendrocyte interface ultrafeatures, such as paranodes, and to provide a platform that links functional electrophysiological enhancements with neuron-glia maturation.

## Conclusions

Human *in vitro* myelination models are essential for dissecting species-specific neuron-oligodendrocyte interactions and enabling mechanistic studies of demyelination with scalable screening of remyelination therapies. While a few multidimensional human systems (e.g., brain organoids) capture some features of neuronal myelination and drug-induced demyelination, they often lack functional readouts that link oligodendrocyte maturation to neuronal activity. To bridge this gap, we developed MyeliMAP, an integrated platform combining PDMS-based soma-exclusion patterning with HD-MEA electrophysiology; to directly couple cellular myelination dynamics to longitudinal functional outcomes. MyeliMAP provides a temporal, quantitative view of how oligodendrocyte maturation shapes neuronal excitability, conduction, and network organization *in vitro*, and offers a human, scalable testbed for probing disease mechanisms and evaluating remyelination strategies across neurodegenerative and demyelinating disorders.

## Supplementary Material

## Supplementary figures

**Figure S1:**
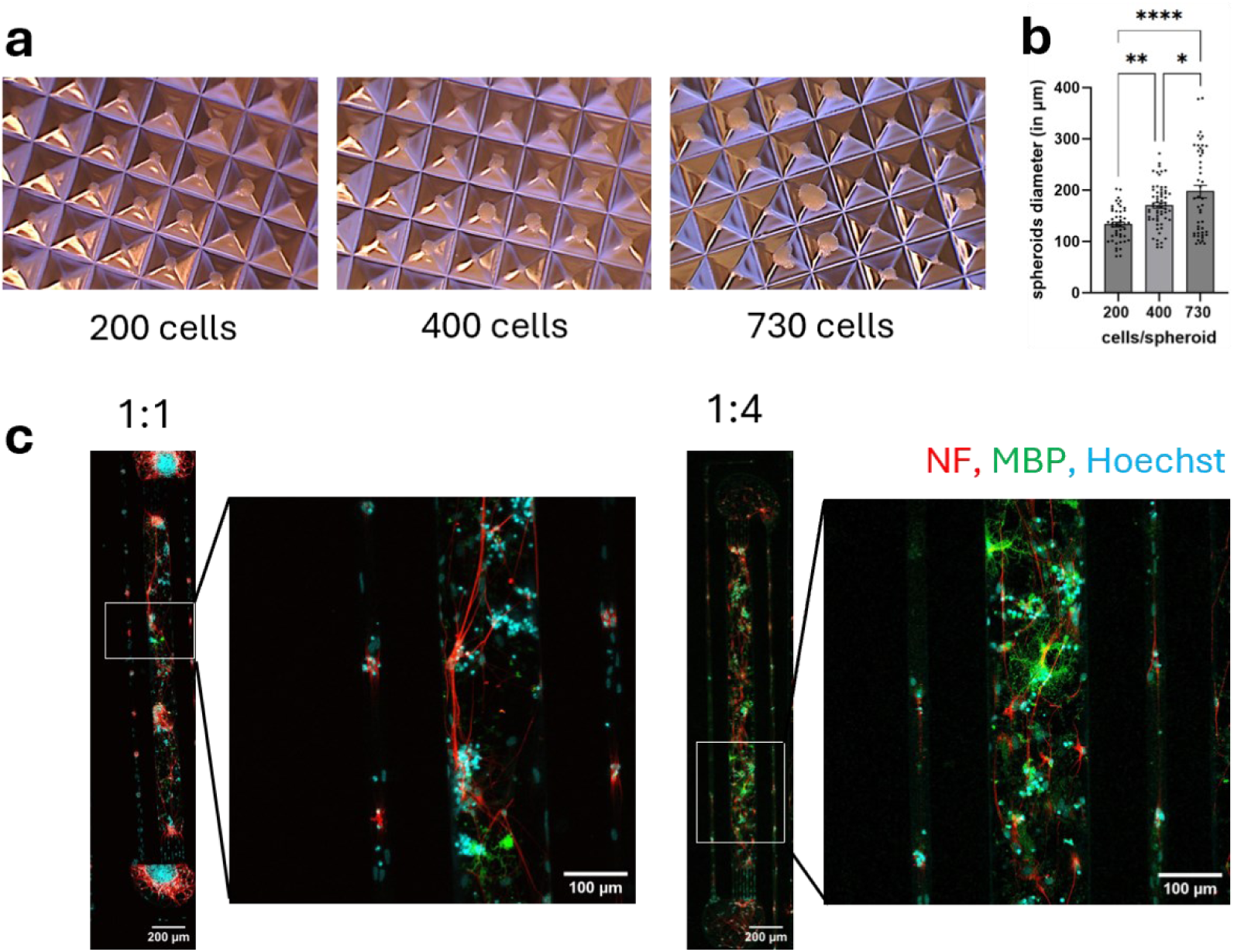
Oligodendrocytes preferentially grow and survive in a soft matrix. **a-b** Brightfield images and quantification of neuronal spheroid diameters generated from 200, 400, and 730 cells per spheroid. **c** Immunofluorescence images of MyeliMAP with 1:1 and 1:4 Matrigel dilutions showing improved oligodendrocyte survival in the softer (1:4) condition. Hoechst (nuclei) images are shown for context. Data shown as mean ± SEM. Statistical analysis was performed by one-way ANOVA with Turkey Multiple Comparison Test (b). *p< 0.05, **p< 0.01, ****p< 0.0001. N = 1 spheroid generation experiment with n=47-63 spheroids (b) and N = 1 MyeliMAP microstructure with n=3 networks (a-c).

**Figure S2:**
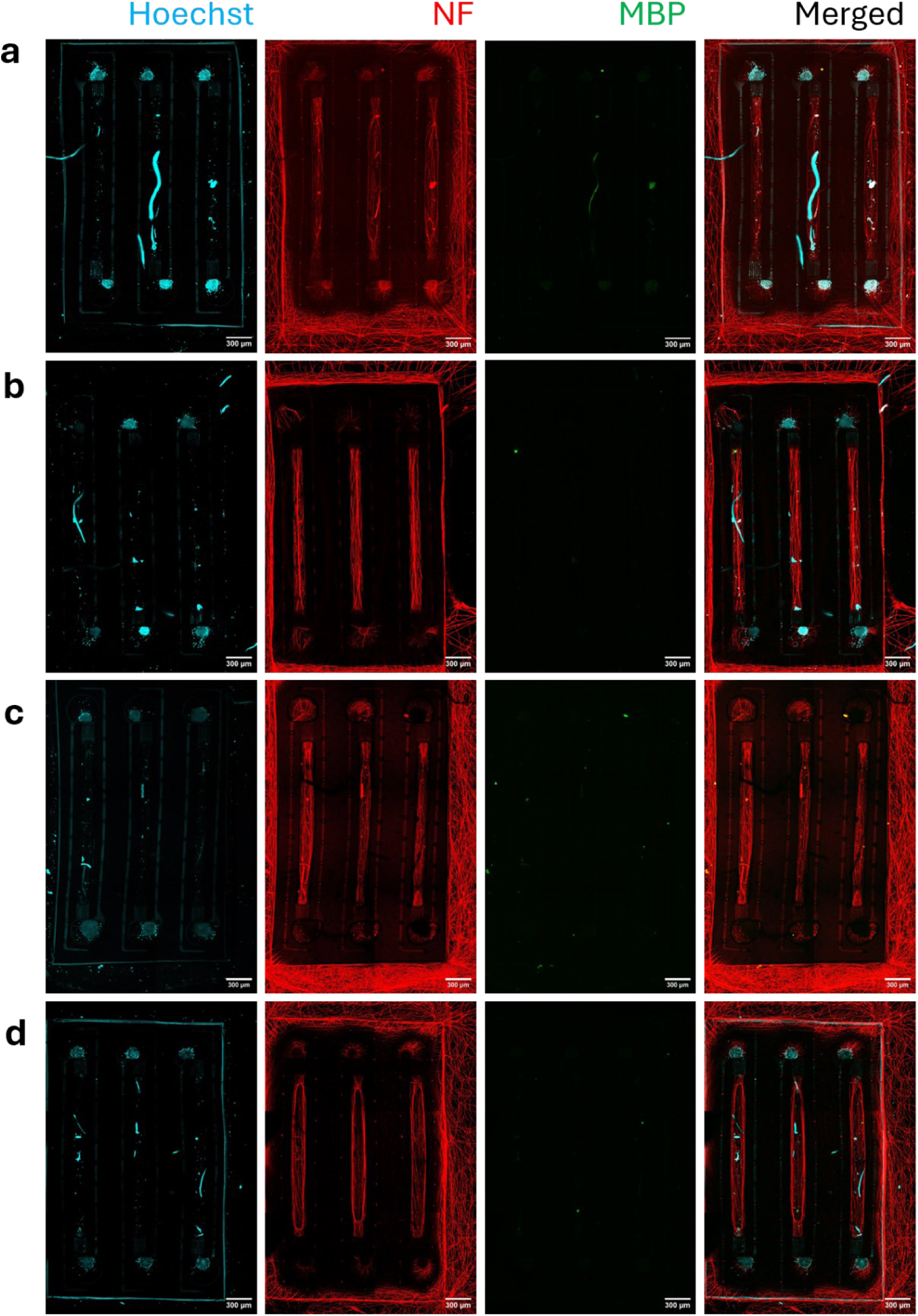
Immunofluorescent staining of MyeliMAPs with neuron monocultures on MyeliMAP-HD-MEA platforms. **a-d** Representative low magnification immunofluorescence images of neuronal monocultures on MyeliMAP-HD-MEA platform after 9 weeks in culture reveals neuronal extensions (NF+) from peripheral seeding wells to central axonal channel where oligodendrocytes (MBP+) are absent. Higher magnification immunofluorescent image from one MyeliMAP-HD-MEA platform (a) showing absence of oligodendrocytes and thus no neuron-oligodendrocyte alignment, shown in Figure 4h. Hoechst (nuclei) images are shown for context. Neuronal extensions are also seen on top and outside of the MyeliMAP microstructure indicating neuronal overgrowth due to presence of 3D extracellular matrix. Scale bar: 300μm. N=4 independent experiments (HD-MEA chips) with n=12 networks.

**Figure S3:**
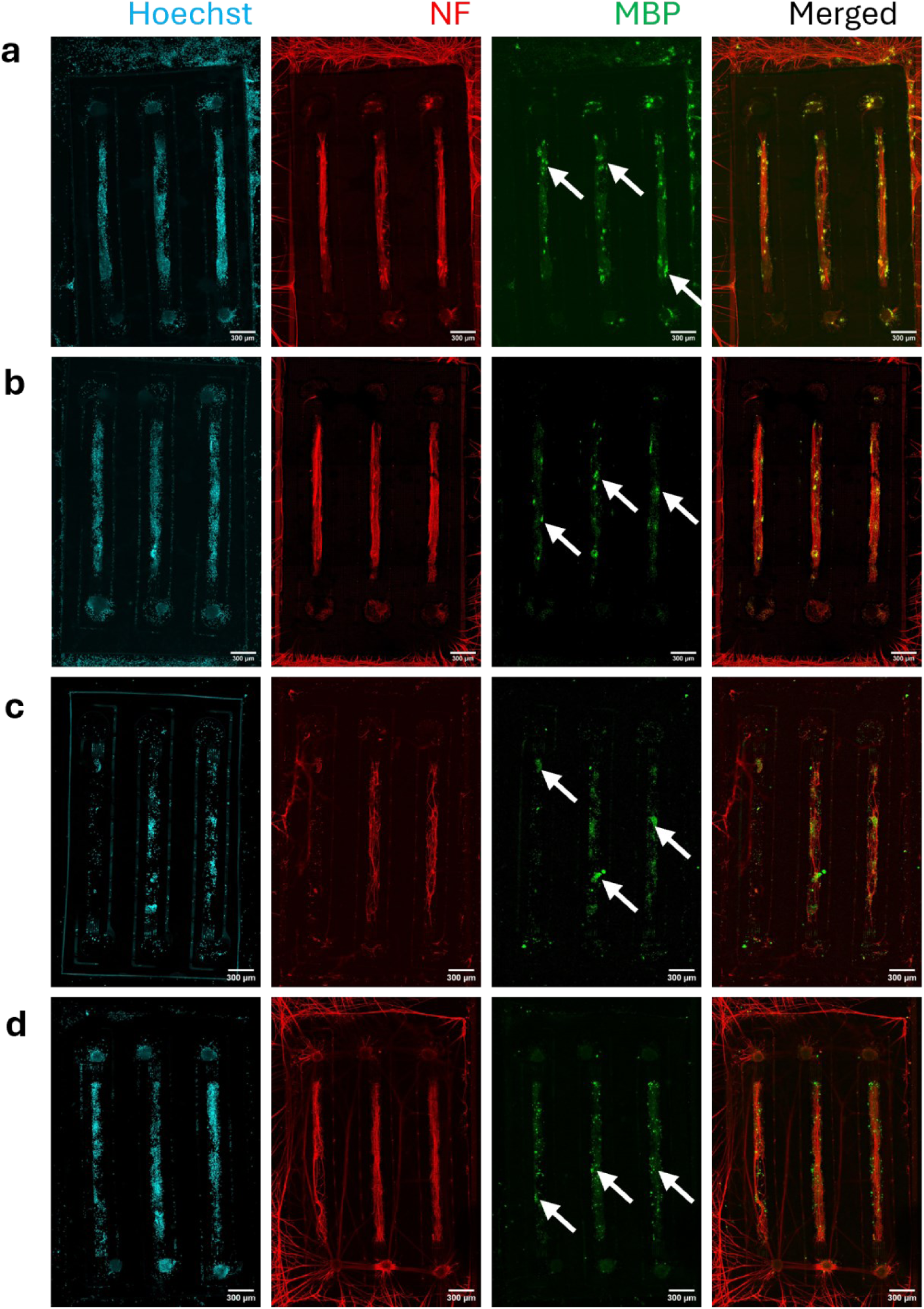
Immunofluorescent staining of MyeliMAPs with neuron-oligodendrocyte cocultures on MyeliMAP-HD-MEA platforms. **a-d** Representative low magnification immunofluorescence images of neuron-oligodendrocyte cocultures grown on MyeliMAP-HD-MEA platforms after 9 weeks in culture reveal neuronal extensions (NF+) from peripheral seeding wells to central axonal channel where they interact with oligodendrocytes (MBP+, indicated with white arrows) to enable neuron myelination. Higher magnification immunofluorescent image from MyeliMAP-HD-MEA platform (a) showing neuron-oligodendrocyte alignment shown in Figure 4h. Hoechst (nuclei) images are shown for context. Neuronal extensions are also seen on top and outside of the MyeliMAP microstructure indicating neuronal overgrowth due to presence of 3D extracellular matrix. Scale bar: 300μm. N=4 independent experiments (HD-MEA chips) with n=12 networks.

## Funding and Acknowledgements

This study was supported by the Research Foundation Flanders (FWO), including individual ongoing fellowships to K.A. (1S03424N), and project grants to Y.C.C., C.V., K.A. (FWO-SBO-S001221N, OrganID; and FWO-G0B5819N). The authors gratefully acknowledge the ETH Zurich-ScopeM facility financing for their support and assistance in this work.

## Contributions

KA designed and performed experiments, analyzed data and wrote the manuscript. BC, AG, JK analyzed the electrophysiology data and contributed in manuscript writing. LM, KW, YCC and CV coordinated the study and contributed in manuscript writing and editing.

## Conflict of interest

The authors declare no conflict of interest.

## Supplementary tables

**Table S1:** Medium composition used in the study.

| Medium | Composition |
| --- | --- |
| **Proliferation Medium** | DMEM/F12 with Glutamax (Lifetech 31331-028), 1x B27 with Vit. A [Life technologies, cat 17504-044], 1x N2 supplements [Invitrogen, cat 17502-048], 10 ng/ml hEGF (Invitrogen, cat PHG0315), 10 ng/ml hFGF (Gibco CTP0263), Pen/Strep (Lifetech 15070-063), Doxycyclin 1ug/ml (Clontech Cat No 631311) |
| **Neuronal Maintenance Medium (NMM)** | NMM comprises of 1:1 mixture of Neurobasal medium (Gibco) and DMEM/F-12 Glutamax™ complimented with 0.5X Glutamax™ (Gibco), 50 U/mL Penicillin-Streptomycin, 0.5X B27 (Gibco), 0.5X N2 (Gibco), 0.5X MEM-NEAA (Gibco), 0.5X sodium pyruvate, 0.025 % human insulin (Sigma) and 50 µM β-mercaptoethonal (Gibco) |
| **Oligodendrocyte Maturation Medium (OMM)** | NMM supplemented with 10 ng/mL insulin growth factor-1 (IGF1), 5 ng/mL hepatocyte growth factor (HGF, Peprotech), 10 ng/mL neurotrophin-3 (NT3, Peprotech), 60 ng/mL triiodothyronine (T3, Sigma), 100 ng/mL Biotin (Sigma), 1 µM cyclic adenosine monophosphate (cAMP, Sigma), and 5 µg/mL doxycycline (Sigma) |
| **Oligodendrocyte Induction Medium (OIM)** | Neurobasal medium supplemented with 0.1 µM retinol acid (RA) (Sigma), 10 µM SB431542, and 1 µM LDN-193189 |
| **Basal medium** | Neurobasal medium supplemented with 10 ng/mL platelet-derived growth factor AA (PDGF-aa, Peprotech), 10 ng/mL insulin growth factor-1 (IGF1), 5 ng/mL hepatocyte growth factor (HGF, Peprotech), 10 ng/mL neurotrophin-3 (NT3, Peprotech), 60 ng/mL triiodothyronine (T3, Sigma), 100 ng/mL Biotin (Sigma), 1 µM cyclic adenosine monophosphate (cAMP, Sigma), and 5 µg/mL doxycycline (Sigma) |

**Table S2:**
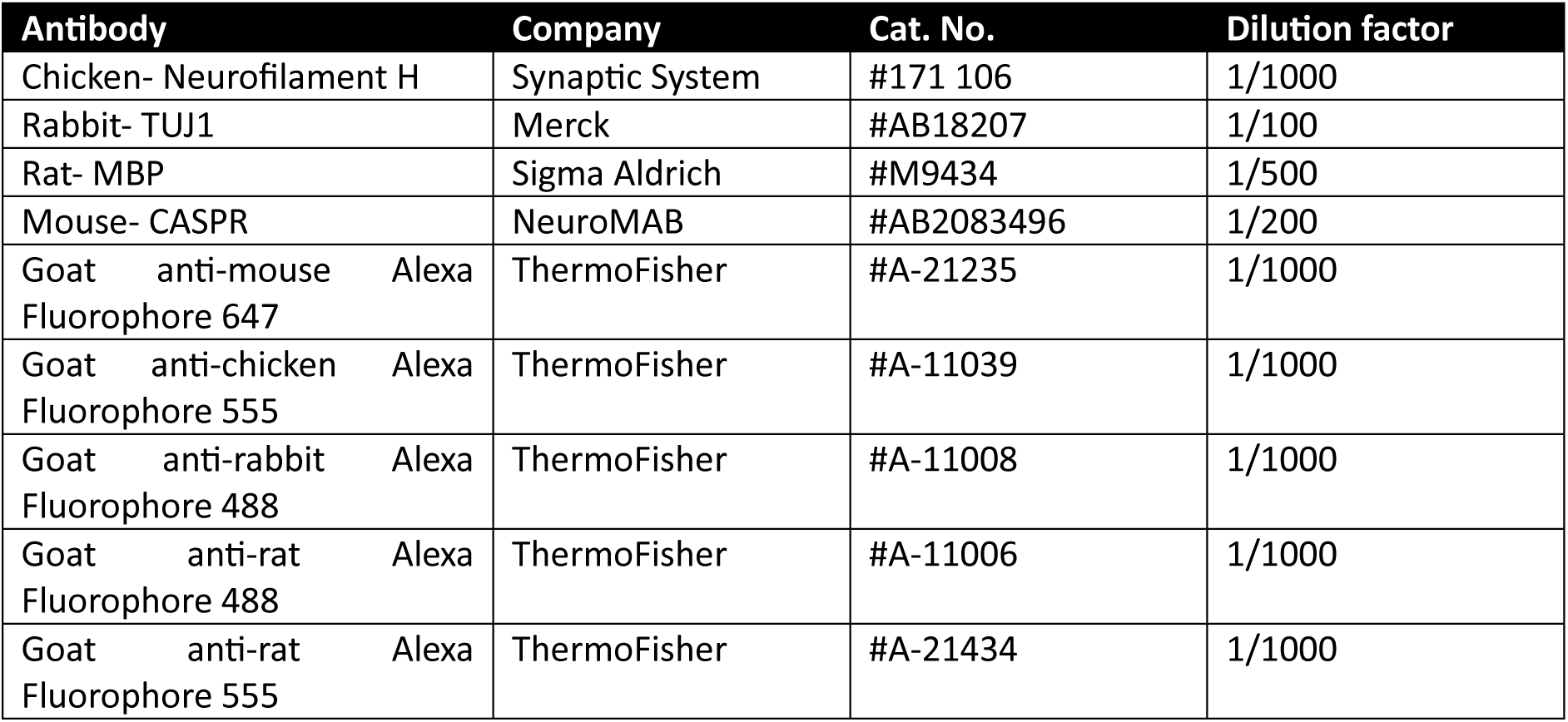
List of primary and secondary antibodies used in the study.

| Antibody | Company | Cat. No. | Dilution factor |
| --- | --- | --- | --- |
| Chicken- Neurofilament H | Synaptic System | #171 106 | 1/1000 |
| Rabbit- TUJ1 | Merck | #AB18207 | 1/100 |
| Rat- MBP | Sigma Aldrich | #M9434 | 1/500 |
| Mouse- CASPR | NeuroMAB | #AB2083496 | 1/200 |
| Goat anti-mouse Alexa Fluorophore 647 | ThermoFisher | #A-21235 | 1/1000 |
| Goat anti-chicken Alexa Fluorophore 555 | ThermoFisher | #A-11039 | 1/1000 |
| Goat anti-rabbit Alexa Fluorophore 488 | ThermoFisher | #A-11008 | 1/1000 |
| Goat anti-rat Alexa Fluorophore 488 | ThermoFisher | #A-11006 | 1/1000 |
| Goat anti-rat Alexa Fluorophore 555 | ThermoFisher | #A-21434 | 1/1000 |

## Notes

### Competing Interest Statement

The authors have declared no competing interest.

